# *f*_4_-statistics-based ancestry profiling and convolutional neural network phenotyping shed new light on the structure of genetic and spike shape diversity in *Aegilops tauschii* Coss

**DOI:** 10.1101/2025.02.16.638466

**Authors:** Yoshihiro Koyama, Mizuki Nasu, Yoshihiro Matsuoka

**Author notes:** Correspondence should be addressed to: Yoshihiro Koyama, Graduate School of Agricultural Science, Kobe University, 1–1 Rokko, Nada, Kobe 657-8501, Japan., Yoshihiro Matsuoka, Graduate School of Agricultural Science, Kobe University, 1–1 Rokko, Nada, Kobe 657-8501, Japan. Abbreviations: CTAB: Cetyltrimethylammonium bromide; CNN: Convolutional neural network; Grad-CAM: Gradient-weighted class activation mapping; MCMC: Markov chain monte carlo; NARO: National Agriculture and Food Research Organization; PCA: Principal component analysis; GRAS-Di: Genotyping random amplicon sequencing-direct; ResNet: Residual neural network; SNP: Single nucleotide polymorphism; XAI: Explainable artificial intelligence.

## Abstract

*Aegilops tauschii* Coss., a progenitor of bread wheat, is an important wild genetic resource for breeding. The species comprises three genetically defined lineages (TauL1, TauL2, and TauL3), each displaying distinctive phenotypes in various agronomic traits, including spike shape. In the present work, we studied the relationship between population structure and spike shape variation patterns using a collection of 249 accessions. *f*_4_-statistics-based ancestry profiling confirmed the previously identified lineages and revealed a genetic component derived from TauL3 in the genomes of some southern Caspian and Transcaucasus TauL1 and TauL2 accessions. Spike shape variation patterns were analyzed using a convolutional neural network-based approach, trained on green and dry spike image datasets. This analysis showed that spike shape diversity is structured according to lineages and demonstrated that the lineages can be distinguished based on spike shape. The implications of these findings for the origins of common wheat and the intraspecific taxonomy of *Ae. tauschii* are discussed.

**Plain Language Summary:** Wild wheat, *Aegilops tauschii*, represents a vast reservoir of alleles that have not yet been utilized in breeding. These alleles may confer beneficial phenotypes, such as drought tolerance and disease resistance, when introduced into bread wheat. To fully leverage this reservoir, it is essential to quickly identify strains with potentially useful alleles. In *Ae. tauschii*, which consists of strain groups (lineages) with unique genetic makeups, this can be done by determining a strain’s lineage based on spike shape. In this work, we trained machine learning models for this purpose and found that spike shape diversity reflects lineage structure. These models demonstrated potential for practical use in assigning strains to their respective lineages based on spike shape. Our work opens new avenues for the application of machine learning in wheat improvement, as well as in the genetic and evolutionary studies of wheat morphology.

## 1. Introduction

*Aegilops tauschii* Coss. (formerly known as *Ae. squarrosa* L.) is a wild annual diploid wheat species native to central Eurasia. Its range extends from Syria and eastern Turkey to Central Asia and China, passing through the Caucasus, northern Iran, Turkmenistan, Afghanistan, and Pakistan. The species’ geographic distribution pattern is distinctive because all other diploid species of the genus *Aegilops* spread westward from the Transcaucasia-northern Iran region. The habitats of this self-pollinating morphologically variable species are ecologically diverse: roadsides, wastelands, arable lands, grasslands, craggy slopes, temperate forests, and sandy seashore.^1)–3)^ Taxonomically, *Ae. tauschii* has two subspecies, *Ae. tauschii* Coss. subspecies *tauschii* and *Ae. tauschii* Coss. subspecies *strangulata* (Eig) Tzvel.^4)^ Subspecies *tauschii* is described as having cylindrical spikes and elongated cylindrical spikelets, whereas subspecies *strangulata* is described as having pearly spikes and thick and bulbous spikelets. Some recent monographs do not formally describe these subspecific taxa, based on the observation that the intermediate forms are often encountered.^1),3)^

*Ae. tauschii* (DD genome) is agronomically important because the species is thought to have given rise to common wheat (*Triticum aestivum* L., AABBDD genome) as the male parent through natural hybridization with *T. turgidum* L. (AABB genome) and its subsequent genome doubling of the F_1_ hybrid.^5–8)^ Studies to understand the *Ae. tauschii*’s natural variation patterns provide valuable basis for wheat improvement because this species serves as a primary genetic resource in breeding practices.^9)^ So far, much work has been done to clarify the genetic structure of *Ae. tauschii.*^10–17)^

Using nuclear and chloroplast DNA polymorphisms, recent population structure studies showed that *Ae. tauschii* has genetically and geographically distinctive lineage structure: two large lineages (TauL1 and TauL2) and a small lineage (TauL3).^18–19)^ TauL1 is widespread within the species range; its sublineage TauL1a occurs mainly in the western part, and TauL1b occurs mainly in the eastern part. TauL2, which is restricted to the western part of the species range with a few exceptions in Xinjiang, China^20)^, has two sublineages: TauL2a, which tends to occur in the western part, and TauL2b, which tend to occur in the eastern part.^21)^ TauL3 is currently known from Georgia only. The genetic distance to the common wheat D genome is shorter in TauL2 and TauL3 than in TauL1. Along with other lines of evidence, this suggests that TauL2 may share a relatively recent ancestor with the common wheat D genome.^18),22)^

No morphological trait that distinguishes those lineages is currently known. In addition, how those lineages relate to the subspecies is not clear. For example, Kihara and Tanaka^23)^ classified only the plants having markedly moniliform (i.e., pearly) spikes as subspecies *strangulata* (i.e., subspecies *strangluata* sensu stricto) and all the rest as subspecies *tauschii* (Fig. 1). This approach had an advantage in unambiguously assigning the intermediate forms to subspecies *tauschii*. However recent studies showed that the subspecies defined using these criteria did not coincide with the major lineages TauL1 and TauL2: subspecies *tauschii* consists of TauL1 and a part of TauL2, whereas subspecies *strangulata* was nested within TauL2.^24)^ In this case, subspecies *strangulata* could be viewed as a form derived from subspecies *tauschii* within TauL2.

**Fig. 1.**
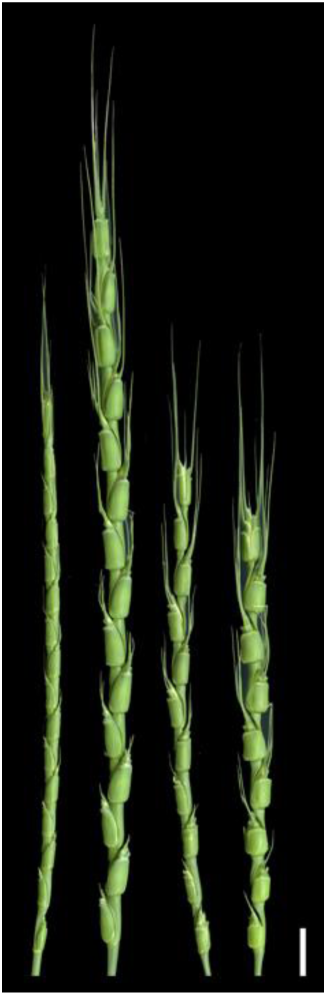
Green spikes of *Ae. tauschii* accessions. A cylindrical spike of IG 48747 (TauL1a) (left), mildly moniliform spikes of KU-20-8 (TauL2a) (center left) and KU-20-10 (TauL2b) (center right), and a markedly moniliform spike of KU-2091 (TauL2b) (right). Of these accessions, subspecies *strangulata* sensu stricto includes only KU-2091, while subspecies *strangulata* sensu lato includes KU-20-8 and KU-20-10, as well as KU-2091. Scale bar: 1 cm.

Another approach to dealing with the intraspecific classification was to include plants having markedly and mildly moniliform spikes in subspecies *strangulata* (subspecies *strangulata* sensu lato) and all the rest as subspecies *tauschii*. These criteria were found useful, because they made subspecies *tauschii* and subspecies *strangulata* largely correspond to TauL1 and TauL2, respectively.^17)^ However, it is unclear to what extent the criteria are usable to classify TauL3. The use of the subspecies index “SI” (glume width/rachis segment width ratio) appears to be in line with these criteria: subspecies *tauschii*, defined by SI, corresponded to TauL1, while subspecies *strangulata* corresponded to TauL2 and TauL3.^25–26)^ Under those criteria, subspecies *tauschii* could be considered as the derived form of subspecies *strangulata.*^27)^

Each *Ae*. *tauschii* lineage displays distinctive phenotypes in various agronomic traits, including flowering time, salt tolerance, and disease resistance.^28–30)^ Thus, methods for correctly assigning *Ae. tauschii* plants to specific lineages based on morphological grounds would be valuable in breeding practice that relies on *Ae. tauschii* as a source of novel alleles. Furthermore, if morphological traits that distinguish the lineages are identified, they could provide avenues for refining the within-species systematics in accordance with the genealogical evidence. To address these issues, it is important to clarify the extent to which spike shape, one of the key traits in the species’ intraspecific classification, distinguishes the *Ae. tauschii* lineages.

In the present work, we studied the relationship between population structure and spike shape variation patterns using a collection of 249 *Ae. tauschii* accessions. Population structure was examined based on single nucleotide polymorphism (SNP) genotypes that were obtained using the genotyping by random amplicon sequencing-direct (GRAS-Di) method.^31)^ Spike shape variation patterns were analyzed using a convolutional neural network (CNN)-based phenotyping method. This method had an advantage in classifying the spike shape continuum between cylindrical and pearly because it can automatically extract spike shape features from digital images and develop classification models based on supervised training that uses the “lineages” (i.e., TauL1, TauL2, and TauL3) as the labels. Furthermore, the explainable artificial intelligence (XAI) algorithms such as Gradient-weighted Class Activation Mapping (Grad-CAM)^32)^ provided visual explanations for the classification models. Therefore, the use of this method had the potential to provide novel insights through computer vision that could complement human vision into how the spike shape differences are associated with the lineage structure.

Our goals in the present study were (1) to clarify the population structure of *Ae. tauschii* based on the genome-wide SNP genotypes of the accessions and discuss its implications for the origins of common wheat and (2) to elucidate the structure of spike shape diversity using convolutional neural network phenotyping and discuss its implications for the intraspecific classification of *Ae. tauschii*.

## 2. Materials and methods

### 2.1. Plant materials

In total, 249 *Ae. tauschii* accessions were used (Table S1). We used 211 accessions representing the entire natural species range for population structure analysis and CNN modeling. The rest 38 accessions, sampled in North Caucasus (Dagestan and Russia), were used for testing the validity of CNN modeling in predicting the lineages based on spike shapes. When the geographical coordinates of the sampling sites were not available in the passport data, we estimated the latitude and longitude using Google Maps (https://maps.google.com/) based on the locality information.

### 2.2. Spike samples

For each accession, well-developed, non-disarticulated spikes were sampled about 14 days after flowering from a single healthy plant grown individually in a pot in a greenhouse during the winter-to-spring season. For 199 accessions, dried spike samples were prepared by air-drying well-developed, non-disarticulated spikes for about six months in envelopes.

### 2.3. SNP genotyping

Total DNA was extracted from the young leaves of a single plant by CTAB (cetyltrimethylammonium bromide) method.^33)^ For each accession, about 10 million raw reads (150 base-pair long, paired-end) were obtained at Eurofins Genomics, Inc (Tokyo, Japan) using the GRAS-Di method.^31)^ The sequencing libraries were prepared by pooling the final product of two sequential polymerase chain reaction steps performed for each accession, using the total DNA as the template and a mix of 12 random primers. The raw reads were generated from the libraries using a NovaSeq 6000 instrument (flow cell type S4) and were subsequently adaptor-trimmed and quality-filtered (by setting the ILLUMINACLIP option to SLIDINGWINDOW:4:30 MINLEN:50) using Trimmomatic.^34)^

Quality reads were aligned to the *Ae. tauschii* AL8/78 reference genome sequence (Aet v5.0)^35)^ using BWA version 0.7.18 (r1243) with the mem option.^36)^ Reads failing to align to the chromosomal sequences were excluded from further analysis. Aligned reads with library insert sizes shorter than 70 base pairs in length or larger than 600 base pairs in length were filtered out using SAMtools version 1.20^37)^, applying the command ‘samtools view-e’((pnext + 150) - pos) >= 70 && ((pnext + 150) - pos) <= 600’’. Duplicated reads were identified and removed using SAMtools. Subsequently, SNPs were called across the accessions relative to the reference genome sequence using BCFtools mpileup version 1.20^38)^ with the setting ‘-q 30’ (minimum mapping quality) and BCFtools call with the setting ‘-G’ (Hardy-Wineberg equilibrium not assumed). The filtering was performed using BCFtools with the setting ‘-i DP>=5 & MQ >= 40’. Sites with a minor allele frequency of less than 5% and missing data frequency exceeding 20% were further filtered out using VCFtools version 0.1.16.^39)^ The resultant SNPs were pruned using plink version 1.90b7, setting the ‘--indep-pairwise’ option to ‘20000 2000 0.5’ for subsequent analyses.^40)^

### 2.4. Principal component analysis

PCA was performed based on a covariance matrix using the probabilistic PCA method available in the pcaMethods (version 1.94.0) package^41)^ for R version 4.4.1.^42)^ The principal component plot was generated using R’s plot function. For spike shape feature vector analysis, a covariance-matrix-based PCA was conducted using the scikit-learn library version 1.5.1 for Python version 3.10.14.^43)^

### 2.5. Struct-f4

We used the Struct-f4 package in the population structure analysis based on the SNP genotypes of the accessions.^44–45)^ Struct-f4 relies on *f*_4_-statistics that estimate amounts of shared drift across pairs of individuals^46)^ and characterizes individual genetic profiles as mixtures of *K* ancestral genetic components without assuming Hardy-Weinberg equilibrium. In the present study, *K* = 5 was chosen, because *Ae. tauschii* had five distinctive lineages/sublineages. ^18–19)^ To estimate parameters, the algorithm was run with 3,000,000 Markov Chain Monte Carlo (MCMC) iterations in the first chain and a burn-in length of 7,500 and then 20,000 MCMC iterations in the second chain.

### 2.6. Geographic distribution map

Maps showing the geographic distribution of the accessions were created based on a spatial dataset obtained from Natural Earth, free vector and raster map data at naturalearthdata.com using the rnaturalearth package.^47)^ The ggplot2 package for R ver. 4.4.1^48)^ was used for plotting.

### 2.7. Spike image data preparation

Spike samples were imaged using a flatbed scanner (EPSON GT-X830). Empty anthers remaining on the fresh spike samples were removed by hand using tweezers prior to imaging. For each accession, fresh or dried spike images (ca. 10 spikes per image) were saved at 96 dpi in a jpg format file with image size equaling to 3400 x 935 pixels.

### 2.8. Supervised learning of residual neural network (ResNet)

A CNN model, ResNet^49)^ was used. ResNet was pretrained on ImageNet-1k^50)^ implemented in PyTorch Image Models library.^51)^ In the green spike image dataset (1,538 images in total), the numbers of images per accession varied from one to nine (mean 7.29), whereas in the dry spike image dataset (2,384 images in total), they ranged from four to 32 (mean 12.04). All images were resized to 400 by 800 pixels (height by width), and their pixel values were normalized using the following parameters: a mean RGB of (0.485, 0.456, 0.406), a standard deviation RGB of (0.229, 0.224, 0.225), and a maximum possible pixel value of 255.0. Supervised training where lineages served as the labels, was conducted using stratified group 5-fold cross-validation. For each green and dry spike image dataset, we split the whole dataset into five equal parts, trained each model on four parts, evaluated it on the fifth, and repeated the process with each part being used as the validation set once. During training, the images were augmented using Albumentations (v1.1.14) library^52)^ with the following transformations: (1) horizontal, vertical, and both horizontal and vertical flips each with a probability of 0.5, (2) random adjustments to brightness and contrast, using default parameters except for brightness_limit, contrast_limit, and p, which were set to 0.2, 0.2, and 0.5, respectively, and (3) random cropping, using default parameters except for height, width, scale, ratio, and p, set to 400, 800, [0.7, 1.0], [2.0, 2.0], and 0.7, respectively. This training pipeline was executed using NVIDIA GeForce RTX 2080 Ti as a hardware accelerator. The macro F1 score, i.e., a harmonic mean of precision and recall over all lineages, was used to evaluate the effectiveness of the trained models in the image classification. The training pipeline was implemented using the PyTorch library version 1.12.1+cu113.^53)^ We used Adam algorithm^54)^ to optimize model parameters based on the value of cross entropy error with learning rate 0.0001. Eight spike images were batched to input into models, and we repeated the training for 60 epochs.

### 2.9. Feature visualization using attention maps

To visualize the image features that influenced the CNN modeling, we used the Grad-CAM, Guided Backpropagation, and Guided Grad-CAM methods to generate attention maps on original spike images based on the final convolutional layer in each ResNet model.^32)^

### 2.10. Spike shape feature vector PCA

The structure of spike shapes diversity was examined by reducing the dimensionality of feature vectors generated in the five ResNet models via PCA for each dry and fresh spike dataset. In each model, feature vectors were obtained for each spike image by conducting a series of convolutions and pooling operations on the input images, with the final full connection layer, which functions as a classifier, being removed. Subsequently, PCA was performed for each model on the averaged feature vectors per accession for dimensionality reduction.

### 2.11. Blind test for lineage prediction

We used 37 of the 38 North Caucasus accessions to test the trained ResNet models for their ability to infer the lineages from spike shape. The image dataset contained 201 green spike images in total and the numbers of images per accession varied from one to eight (mean 5.43). All images were resized to 400 by 800 pixels (height by width), with pixel values normalized according to these parameters: mean RGB values of (0.485, 0.456, 0.406), standard deviation RGB values of (0.229, 0.224, 0.225), and a maximum possible pixel value of 255.0. The lineage membership probabilities for each accession were calculated by feeding the image dataset to each of the five ResNet models trained on the green or dry spike images of the 211 accessions. In each spike image dataset, the resultant probabilities were averaged for each lineage, with the lineage exhibiting the highest probability inferred as the one to which the accession belonged. Concurrently, the 38 North Caucasus accessions underwent SNP genotyping using the GRAS-Di method. SNPs were called as described for the 211 accessions used for the ResNet modeling. The merged raw SNPs (called for 249 accessions in total) were then filtered and pruned as described. Subsequently, each North Caucasus accession was assigned to a lineage based on the result of a PCA performed on the SNP genotypes. The accuracy of the spike-image-based inference for lineage was assessed by comparing it to the genotypic assignment for consistency.

## 3. Results

### 3.1. SNP genotyping

In total, qualified SNPs at 16,746 sites across the 211 accessions relative to the AL8/78 reference genome sequence were obtained. The average observed heterozygosity varied from 0.06 to 0.11 between accessions (mean 0.09). The AL8/78 accession in our panel shared the same nucleotide with the reference genome sequence at 12,399 out of 13,936 homologous sites (89.0%). The SNP sites were distributed widely over the genome (Fig. 2A). For each chromosome, numbers of the sites were 1,988 for chromosome 1D, 2,951 for 2D, 2,621 for 3D, 1,898 for 4D, 2,666 for 5D, 2,062 for 6D, and 2,560 for 7D.

**Fig. 2.**
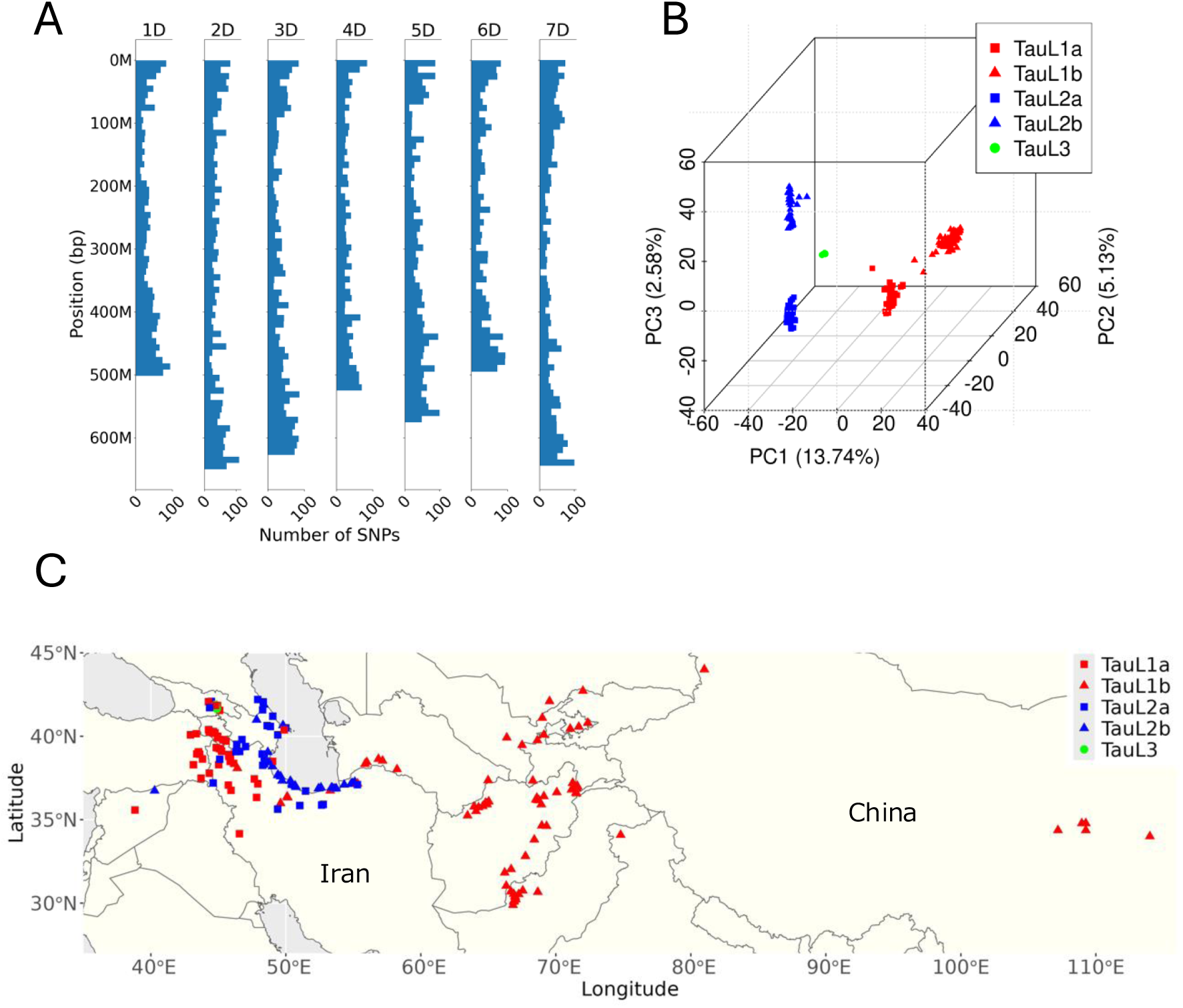
GRAS-Di-derived SNP genotyping of the 211 accessions. (A) Distribution of GRAS-Di-derived SNPs mapped to the *Ae. tauschii* AL8/78 reference genome sequence. Bar plots of the number of markers are displayed every 10 Mbp. (B) Graph of the first three axes principal components from a PCA based on the GRAS-Di-derived SNP genotypes of the 211 accessions. The first component (PC1) accounts for 13.74%, the second (PC2) for 5.13%, and the third (PC3) for 2.58% of the total variance. The lineage and sublineage are represented by color and shape for each accession according to the key. (C) Geographic distribution of the 211 accessions. The lineage and sublineage are represented by color and shape for each accession according to the key.

### 3.2. Population structure

A PCA of the SNP genotypes revealed that the 211 accessions were categorized into the known lineages/sublineages: TauL1 (consisting of TauL1a and TauL1b), TauL2 (consisting of TauL2a and TauL2b), and TauL3 (Fig. 2B). Two exceptions were noted: KU-2109 (TauL1, previously classified as TauL2) and IG 127015 (TauL2, previously classified as TauL1).^18)^ TauL1 (135 accessions) was spread across the species range, whereas TauL2 (71 accessions) was exclusively located in the western part of the range. TauL3 (five accessions) was confined to Georgia (Fig. 2C). Moreover, within the TauL1 lineage, the sublineage TauL1a (52 accessions) predominantly occurred in the western region, while TauL1b (83 accessions) was primarily observed in the eastern part. Similarly, the sublineage TauL2a (35 accessions) tended to occur in the western region of the lineage range, whereas TauL2b (36 accessions) showed a preference for the eastern part (Fig. 2C). All these observations aligned with prior research findings.^18),21)^

To further examine the population structure and genetic profiles of individual accessions, a Struct-f4 analysis (*K* = 5) was conducted. This analysis showed that, within each lineage/sublineage, the accessions shared a specific ancestral genetic component that served as a major factor in their genetic profiles (Fig. 3A; Table S1). The proportions of these components were greater than 0.75 in each accession, with a few exceptions (KU-2109 in TauL1a, KU-2068, KU-2122, and KU-2153 in TauL1b, KU-2158 in TauL2b), supporting the PCA observations that each lineage/sublineage is a distinctive genetic aggregate. Within the TauL1 and TauL2 lineages, these components were shared as minor factors by many accessions between TauL1a and TauL1b and between TauL2a and TauL2b. In contrast, the proportions of these components in the genetic profiles were negligible in most accessions belonging to different lineages (i.e., TauL1 vs. TauL2). These observations provided evidence that, despite the clear separation between the sublineages observed in the PCA plots, TauL1a and TauL1b, and TauL2a and TauL2b, form distinct lineages, TauL1 and TauL2, respectively (Fig. 3A).

**Fig. 3.**
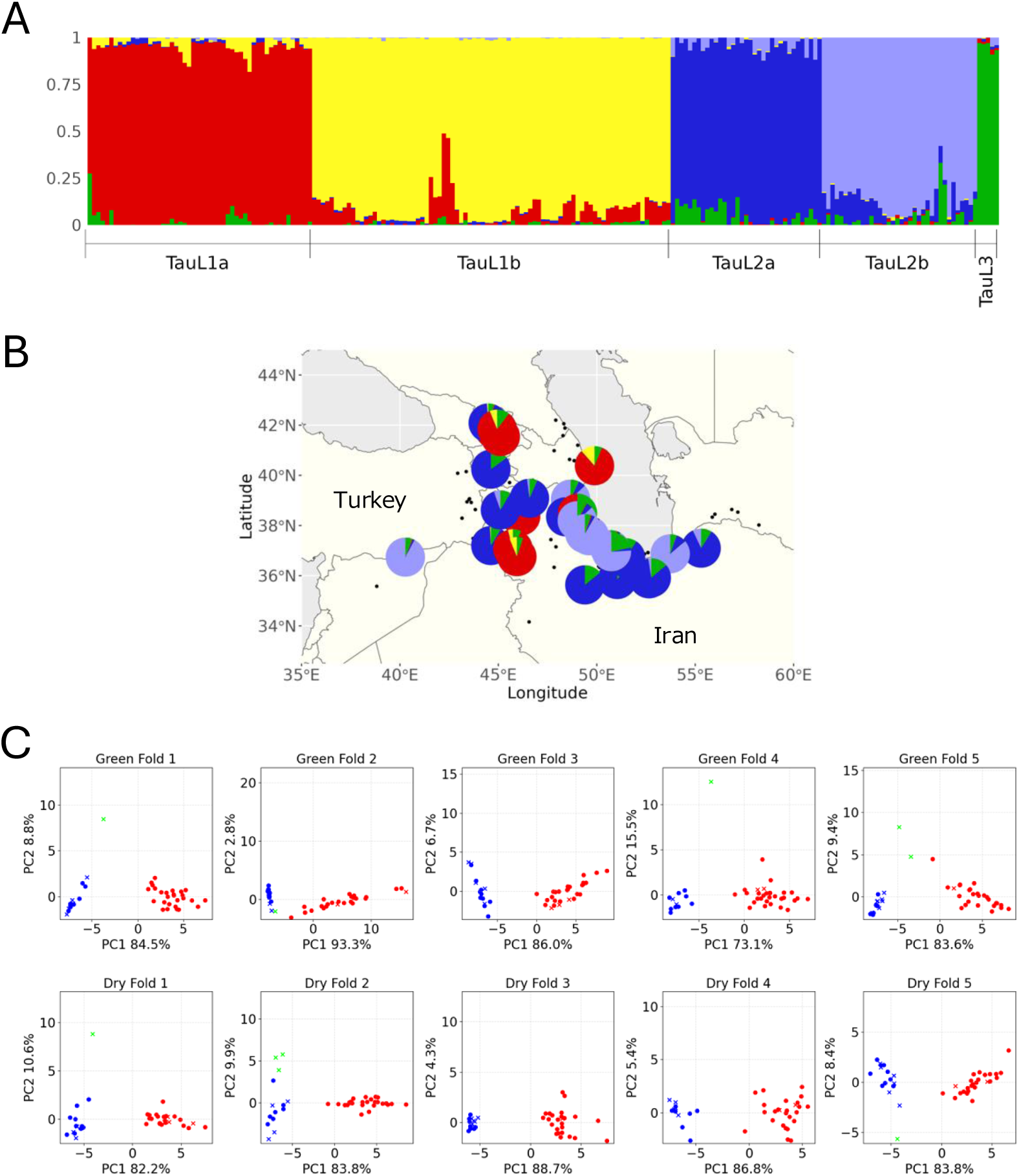
*f*_4_-statistics-based ancestry profiling and convolutional neural network spike shape phenotyping in the 211 accessions. (A) Proportions of the ancestral genetic components of the 211 accessions based on *f*_4_-statistics-based ancestry profiling at *K* = 5. The ancestral genetic components for TauL1a, TauL1b, TauL2a, TauL2b, and TauL3 are represented in red, yellow, blue, purple, and green, respectively. (B) Geographic distribution of *Ae. tauschii* accessions with TauL3 ancestral component proportions greater than 0.05, represented by pie charts. The ancestral genetic components are color-coded as in (A). Black points denote accessions with TauL3 ancestral component proportions of 0.05 or less. (C) Graphs of the first two axes (PC1 and PC2) from PCAs performed on averaged feature vectors per accession for trained ResNet models, generated using the green and dry spike image validation datasets in 5-fold cross-validation. The variance explained by each axis is shown in each graph. The lineages are color-coded: red for TauL1, blue for TauL2, and green for TauL3. Crosses denote accessions with TauL3 ancestral component proportions greater than 0.05.

This analysis further indicated that certain TauL1 and TauL2 accessions exhibited small but substantial proportions of the genetic component originating from TauL3. The proportions of the TauL3 component ranged from 0.00 to 0.27 (mean 0.02) in TauL1a, from 0.00 to 0.05 (mean 0.01) in TauL1b, from 0.00 to 0.15 (mean 0.05) in TauL2a, from 0.00 to 0.33 (mean 0.04) in TauL2b, and from 0.91 to 0.97 (mean 0.95) in TauL3. The TauL1 and TauL2 accessions with proportions greater than 0.05 (33 accessions in total) were confined to TauL1a (eight accessions), TauL2a (15 accessions), and TauL2b (10 accessions), and geographically limited to the coastal Caspian as the center, with the Transcaucasus and adjacent regions (Fig. 3B).

### 3.3. Conventional neural network analysis of spike shape

We analyzed the natural variation patterns in *Ae. tauschii* spike shape based on CNN, aiming to clarify how its diversity is structured in relation to the population structure of the species. Firstly, we trained ResNet models^49)^ using the lineages as the labels to predict the lineage (TauL1, TauL2, and TauL3) to which an accession belongs based on its spike shape. The 5-fold cross-validation generated five trained ResNet models for each green (1,538 images in total) and dry (2,384 images in total) spike image dataset. The trained ResNet models demonstrated very high macro F1 scores (0.95 – 1.00), indicating their highly effective assignment of the spike images to the correct lineages (Table 1). Next, PCA was performed on averaged feature vectors per accession for each trained ResNet model. For the green and dry spike image datasets, TauL1 and TauL2 formed separate clusters on the PC1 axis in each PC plot (Fig. 3C). This indicated that the spike shapes of the TauL1 and TauL2 accessions were similar within lineages compared to between lineages. When included in the fold, TauL3 tended to be separate from TauL1 and TauL2 on the PC2 axis. In each plot, accessions with the TauL3 component proportions greater than 0.05 showed no clear distribution pattern within their respective lineages (Fig. 3C).

**Table 1.** Macro F1 scores of the five trained ResNet models, generated by stratified group 5-fold cross-validation, for the green and dry spike image datasets.

| Dataset | Model1<br>(fold1) | Model2<br>(fold2) | Model3<br>(fold3) | Model4<br>(fold4) | Model4<br>(fold5) |
| --- | --- | --- | --- | --- | --- |
| Green spike image | 1 | 0.98 | 0.99 | 0.99 | 0.96 |
| Dry spike image | 0.98 | 0.95 | 1 | 0.99 | 0.99 |

The trained ResNet models were further tested for their ability to predict the lineage to which an accession belongs based on its spike shape. Of the 38 North Caucasus accessions with unknown lineages, green spike images, which were obtained for 37 accessions (201 images in total), were used in the test. The ResNet models trained on green spike images classified the accessions into TauL1 (19 accessions) and TauL2 (18 accessions) based on their spike shapes (Table S2). Interestingly, the ResNet models trained on dry spike images produced the same classification, assigning the same 19 accessions to TauL1 and the remaining 18 to TauL2 (Table S2). This CNN classification was fully consistent with the categorization based on the genotypes: PCA of the genotypes based on 15,447 GRAS-Di-derived SNPs showed that each North Caucasus accession belonged to TauL1a or TauL2a (Fig. 4).

**Fig. 4.**
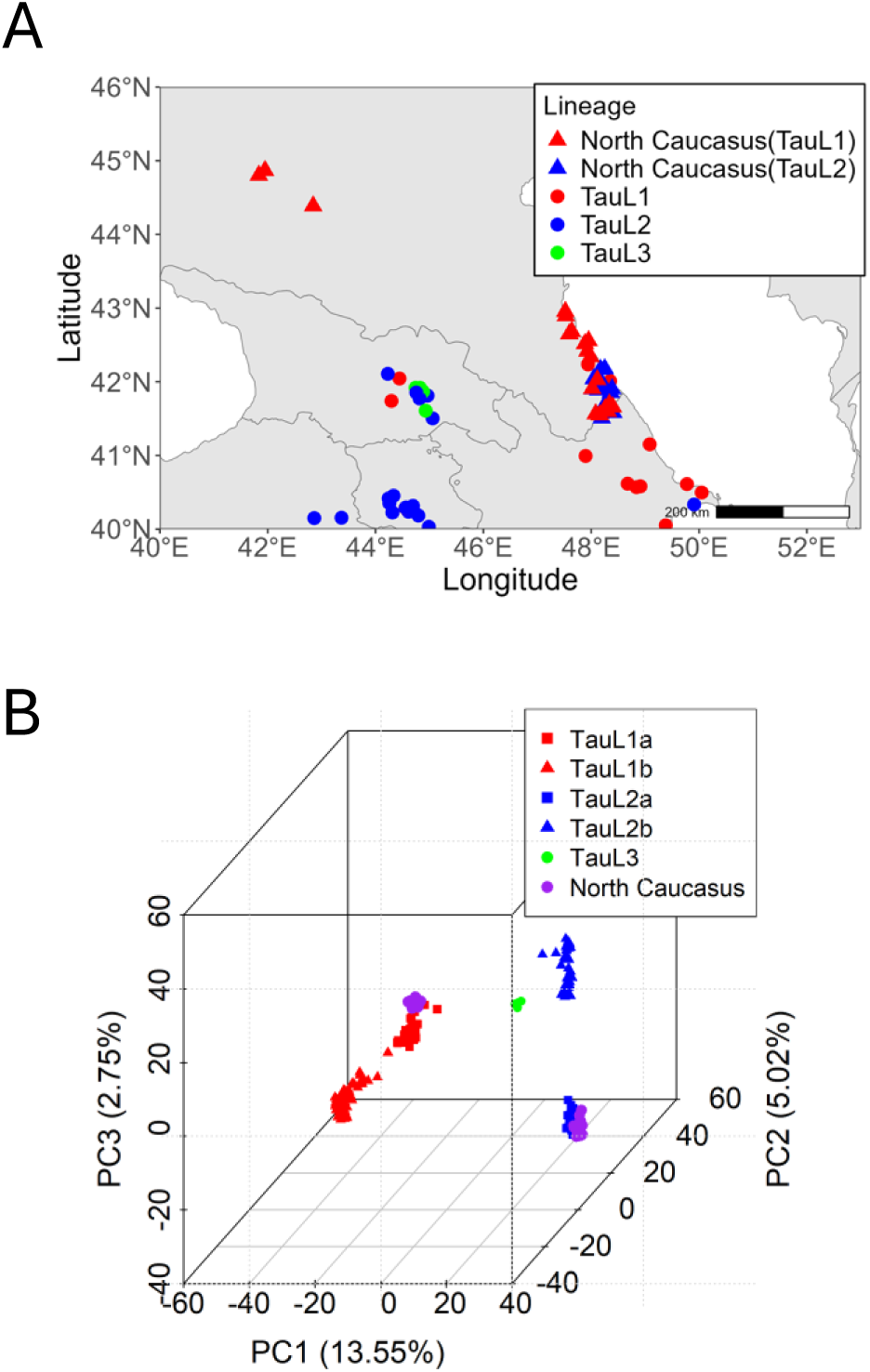
Population structure and geographic distribution and of the Caucasus accessions. (A) Graph of the first three principal components from a PCA based on the GRAS-Di-derived SNP genotypes of 249 accessions, including the 38 North Caucasus accessions used for the blind test. The first component (PC1) accounts for 13.55%, the second (PC2) for 5.02%, and the third (PC3) for 2.75% of the total variance. The lineage and sublineage are represented by color and shape for each accession according to the key. (B) Geographic distribution of *Ae. tauschii* accessions in the Caucasus region. North Caucasus accessions used for the blind test are represented by triangles, while other accessions are represented by circles. The lineage and sublineage are color-coded for each accession according to the key.

In both green and dry spike image datasets, the image features influencing the ResNet modeling varied across different spike images within individual accessions, regardless of their respective lineages. For example, in the case of the dry spike images of KU-2091 (TauL2, 14 images in fold1), the features were observed at the tips in some attention map images, whereas they were distributed across the entire spikes in others (Fig. 5). Similarly, in the case of the green spike images of IG 48748 (TauL1, eight images in fold2), the features localized in the central parts in some attention map images, whereas they showed a broad distribution across the central spikes in others (Fig. 5).

**Fig. 5.**
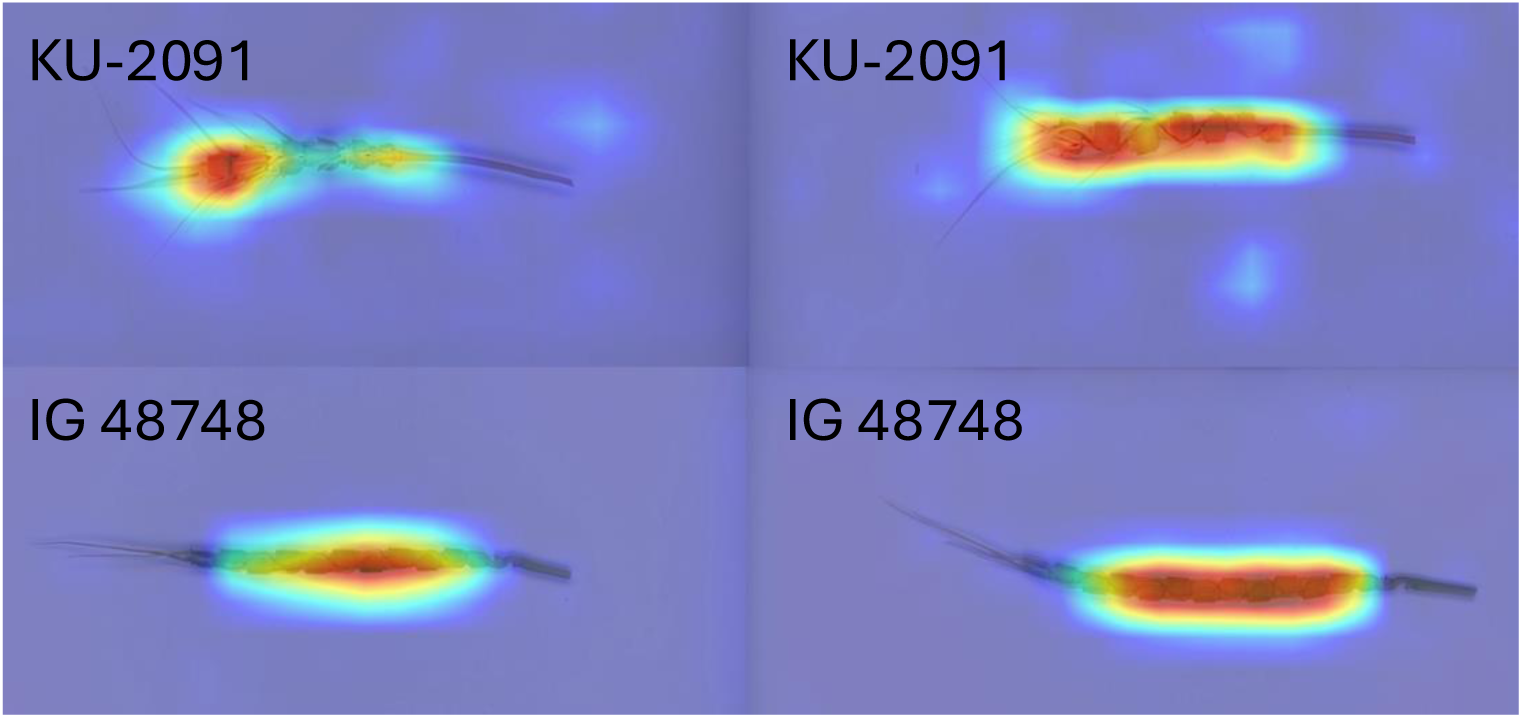
Examples of spike shape attention map images generated using the Grad-CAM method. The color gradient ranges from high attention (red, orange, and yellow) to low attention (green and blue). The *Ae. tauschii* accession from which the spike is derived is shown in each figure. All spike shape attention map images generated using the Grad-CAM, Guided Backpropagation and Guided Grad-CAM methods are available in the Zenodo repository.

## 4. Discussion

### 4.1. Population structure of *Ae. tauschii* and its implications for the origins of common wheat

Based on the PCA and Struct-f4 analyses of GRAS-Di SNP genotypes of the accessions, the present study showed that the *Ae. tauschii* population comprises three genetically and geographically distinctive lineages/sublineages: TauL1 comprising TauL1a and TauL1b, TauL2 comprising TauL2a and TauL2b, and TauL3. This finding aligns with previous research.^18),19)^ In the lineage assignment, two accessions were placed in the lineages different from those to which they had previously been assigned based on Diversity Arrays Technology marker genotypes. KU-2109, formerly classified as TauL2, was reassigned to TauL1, while IG 127015, formerly classified as TauL1, was reassigned to TauL2.^18)^ The reassignment of KU-2109 (collected in Iran) to TauL1 is consistent with a recent study that used *k*-mers for genotyping.^55)^ Further analysis is required to confirm the correct lineage of IG 127015 (collected in Armenia); however, for the purpose of this study, the accession was included in TauL2.

In the Struct-f4 analysis, several TauL1 and TauL2 accessions were found to have non-negligible proportions of the genetic component originating from TauL3 (Fig. 3B). Gene flow from TauL3 in ex-situ genebank fields, where the materials were propagated, might partially explain the sharing of the TauL3 genetic component, as its relatively long anthers are capable of shedding pollen.^56)^ However, it is more likely that the observed pattern of TauL3 component sharing originated from inter-lineage hybridization in natural habitats, rather than cross-pollen contamination in modern ex-situ fields. This conclusion is based on the following considerations. First, the TauL3 component was restricted to the southern Caspian and Transcaucasus. This geographic distribution pattern contrasts with the random distribution expected if gene flow had occurred in the ex-situ genebank fields. Second, 26 out of 33 accessions with TauL3 genetic component proportions greater than 0.05 (Table S1), including KU-2109 (TauL1a, Iran), KU-2158 (TauL2b, Iran), and KU-2159 (TauL2b, Iran), which exhibit exceptionally high TauL3 genetic component proportions (> 0.2), were provided by the National Bioresource Project/Kyoto University. These materials have been maintained through selfing via spike-bagging to prevent cross-pollen contamination since the introduction of the original samples collected from their respective natural habitats. Therefore, admixture within and between lineages may have shaped the observed distribution pattern of the TauL3 component. TauL1 and TauL2 accessions with TauL3 genetic component proportions greater than 0.05 were found primarily along the coastal Caspian, as well as in the Transcaucasus and adjacent regions. This may suggest that these areas served as centers of admixture involving this component (Fig. 3B).

Regarding the TauL3 component in the D genome, common wheat cultivars can be categorized into two types: those with little to low proportions of the TauL3 component, distributed across Eurasia and northern Africa, and those with substantial proportions, largely confined to the Transcaucasus and adjacent regions, including Georgia, Armenia, and the border areas of Turkey, Iraq, and Iran.^55)^ Importantly, more than 70% of the common wheat D genome exhibits “identity-by-state” with the genomes of southern Caspian TauL2 sublineages.^55)^ However, because common wheat D genome consistently forms a distinct cluster from *Ae. tauschii* in population structure studies^12),14),18)^, the direct descendants of the *Ae. tauschii* populations that gave rise to the common wheat D genome have likely not yet been identified.

The admixed nature of *Ae. tauschii* genomes, along with these previous findings, may have important implications for the evolution of common wheat. It can be inferred that the first type of cultivars likely originated from hybridizations involving *Ae. tauschii* populations with low proportions of the TauL3 component. However, those ancestral *Ae. tauschii* populations may not have been the direct progenitors of the current southern Caspian TauL2, although they were genetically closely related to it, as suggested by the findings mentioned above. The second type of cultivars likely originated from hybridizations involving TauL2-related *Ae. tauschii* populations that contained substantial TauL3 component proportions. A caveat to this non-exclusive scenario is the absence of populations genetically closely related to the current southern Caspian TauL2 (TauL2b) in the centers of the second type of cultivars, the Transcaucasus and adjacent regions (Fig. 3B). Since little is known about the historical genetic changes experienced by *Ae. tauschii* populations, addressing this knowledge gap may help further clarify the origins of common wheat.

### 4.2. Structure of spike shape diversity and its implications for intraspecific taxonomy

The common wheat D genome inherited only a small portion of *Ae. tauschii* alleles due to the limited number of ancestral populations (i.e., those closely related to the current southern Caspian TauL2 and Georgian TauL3) involved in natural hybridizations with *T. turgidum*. Consequently, modern *Ae. tauschii* represents a vast reservoir of alleles that have not yet been utilized in breeding. Such alleles may confer beneficial phenotypes, such as drought tolerance and disease resistance, when introduced into common wheat. In this context, TauL1, the most distant relative of the common wheat D genome, stands out as a rich source of such alleles, while each lineage could provide valuable genetic resources for wheat breeding. To efficiently utilize *Ae. tauschii* germplasm, it is essential to develop methods for distinguishing the lineages based on accession morphology, without the need for genotyping.

*Ae. tauschii* exhibits considerable diversity in spike shape, with some accessions having cylindrical outlines, others curved, and a range of intermediate forms. In the present study, we aimed to clarify the structure of spike shape diversity using CNN modeling. We found that, in the 249 accessions used, the lineages can be distinguished based on spike shape. Notably, the CNN-based approach demonstrated impressive potential for practical use in assigning accessions to the correct lineages based on the spike images. This was evident from the very high macro F1 scores (0.95 – 1.00) obtained for the trained ResNet models (Table 1), the clear, non-overlapping separation of the lineages on the feature vector PC plots (Fig. 3C), and the fully accurate assignment of the 37 North Caucasus accessions to their respective lineages in the blind test (Fig. 4A; Table S2). In the blind test, the ResNet models trained on dry spike images classified the green spike images into the correct lineages. This indicates that the models rely on the spike shapes rather than the colors when making classifications. Nevertheless, we caution that the under-representation of TauL3 accessions in our samples due to limited availability may limit the ability of the CNN-based approach to distinguish this lineage from TauL1 and TauL2. The usefulness of this method for TauL3 must therefore be critically re-evaluated.

A taxonomy should provide a useful tool for field biologists while reflecting the phylogenetic history of the taxa as accurately as possible.^57)^ Considering this perspective, our findings have implications for the intraspecific taxonomy of *Ae. tauschii*. First, TauL1, TauL2, and TauL3 should each ultimately be treated as a distinct taxon, as each represents a well-defined genealogical unit. This may require reconsidering the dichotomous nature of the current system—characterized by cylindrical spikes and elongated cylindrical spikelets in subspecies *tauschii*, and pearly spikes and thick, bulbous spikelets in subspecies *strangulata*—because it does not explicitly account for the existence of three distinct lineages. The sublineages of TauL1 and TauL2 could also be ranked as taxa within their respective lineage-level taxa. Second, to achieve this goal, morphological keys to the lineages and sublineages—essential for field biologists and breeders engaged in classification—must be identified. Spike shape remains the preferred trait for this purpose, as the ResNet models demonstrated its potential for distinguishing the lineages. In the present work, we were unable to identify such key phenotypes from the spike image features influencing the ResNet modeling, as visualized on the attention maps (Fig. 5). Further examination through human observation, combined with CNN-based analyses using a large spike image dataset that includes more representation of the TauL3 lineage, may help address this issue.

## Acknowledgements

We thank Pablo Librado for his assistance with running Struct-f4. We also extend our gratitude to Kazutoshi Okuno for providing collection site information on the 38 North Caucasus accessions, to the Genebank project at the National Agriculture and Food Research Organization (NARO) for supplying the seeds of these accessions, and to Hirokazu Handa for providing the seeds of AL8/78. Computations were partially performed on the NIG supercomputer at ROIS National Institute of Genetics. This work was supported by JPSP KAKENHI Grant (19H02935, 19KK0157, 20K20720, 23K23573, and 23K26927), Moonshot Agriculture, Forestry and Fisheries Research and Development Program (JPJ009237), Hyogo Science and Technology Association Academic Research Grant (number 4002, fiscal year 2022), and the Joint Research Program of Arid Land Research Center, Tottori University (04A2002 and 06B2007).

## Supplemental Materials

Table S1. The *Ae. tauschii* accessions used and their sources, geographic properties, and genetic properties.

Table S2. The lineage membership probabilities and predicted lineages obtained for the 37 North Caucasus accessions in the blind test, along with their sublineages defined based on GRAS-di-derived SNP genotypes.

Other datasets and source codes generated during and/or analyzed during the current study are available in the DDBJ BioProject repository accession number PRJDB19942: https://ddbj.nig.ac.jp/search/entry/bioproject/PRJDB19942 and the Zenodo repository:

Koyama, Y., Nasu, M., & Matsuoka, Y. (2025). Data from: f4-statistics-based ancestry profiling and convolutional neural network phenotyping shed new light on the structure of genetic and spike shape diversity in Aegilops tauschii Coss. [Data set]. Zenodo. https://doi.org/10.5281/zenodo.14627552

Koyama, Y. (2025). yox5ro/ae-tauschii-lineage-classifier-cnn: Initial release (1.0.0). Zenodo. https://doi.org/10.5281/zenodo.14697946

